# FSL-CP: A Benchmark for Small Molecule Activity Few-Shot Prediction using Cell Microscopy Images

**DOI:** 10.1101/2023.10.11.560835

**Authors:** Son V. Ha, Lucas Leuschner, Paul Czodrowski

## Abstract

Predicting small molecule activities using information from high-throughput microscopy images has been shown to tremendously increase hit rates and chemical diversity of the hits in previous drug discovery projects. However, due to high cost of acquiring data or ethical reasons, data sparsity remains a big challenge in drug discovery. This opens up the opportunity for few-shot prediction: fine-tuning a model on a low-data assay of interest after pretraining on other more populated assays. Previous efforts have been made to establish a benchmark for few-shot learning of molecules based on molecular structure. With cell images as a molecular representation, methods in the computer vision domain are also applicable for activity prediction. In this paper, we make two contributions: a) A public data set for few-shot learning with cell microscopy images for the scientific community, b) A range of baseline models encompassing different existing single-task, multi-task and meta-learning approaches.

## Introduction

High-throughput Imaging (HTI) has been a powerful tool in drug discovery, having yielded many biological discoveries.^1–3^ It often involves capturing the morphological changes of the cells induced by chemical compounds, and quantifying these changes into a large set of numerical features^4^ such as staining intensity, texture, shape ans spatial correlations. They act as ‘fingerprints’ that can be used to characterise compounds in a relatively unbiased way. This technique, known as morphological profiling, has proven to be useful for a variety of applications, such as optimizing the diversity of compound libraries, ^5^ determining mechanism of actions of compounds, ^6–8^ and clustering of genes by their biological functions. ^9,10^

### Small Molecule Activity Prediction

Prediction of small molecule activity against a drug target is an important task in drug discovery. It helps identify and optimise compounds for a desired activity, as well as recognise and avoid off-target activities. This leads to the identification of the compounds with highest potential in the early drug discovery pipeline.

HTI data has been used in bioactivity prediction by Simm et al.^11^ in two drug discovery projects, which lead to a tremendous increase in hit rates by 50- to 250-fold, while increasing the chemical structure diversity of the hits. In these projects, only 1.6% of the label matrix is filled, for over 500,000 compounds and 1200 prediction tasks. This reflects the need for a modeling paradigm which not only can adapt to new tasks quickly with little data, but also leverage the availability of many low-data related tasks. We find this setting is ideal to form a *few-shot learning* challenge.

### Few-shot Learning

In the few-shot learning setting, there is no one big dataset *D* to learn from, but instead many small datasets we called *tasks*, denoted *T* . The aim of few-shot methods is to generalise over new tasks 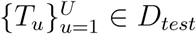 efficiently with only a small number of available datapoints. Each task *T*_*u*_ consists of a *support set S* for learning, and a *query set Q* for evaluation, *T*_*u*_ = ⟨*S, Q*⟩. Typically the size of support set *S* is very small to reflect the low-data setting.

Few-shot models adapt efficiently to low-data tasks by using an advantage initialisation of its parameters, normally though some sort of *pretraining* on a large data corpus such as a set of auxiliary tasks 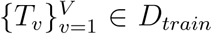. We expect that knowledge gained from pretraining can be transferred effectively to new unseen tasks, so that models can quickly learn these new tasks using only little data. This can be compared to, for example, a person who already has prior knowledge of music can pick up a new musical instrument relatively fast with little demonstration.

Most state-of-the-art few-shot methods come from computer vision and natural language processing domain.^12–16^ Drug discovery is another field where there is growing interest in few-shot learning,^17^ since data scarcity is a common setting for many prediction tasks. In this paper, we propose to expand another challenge in drug discovery to the few-shot learning area: the aforementioned small molecule activity prediction with cell imaging data. We find that this is a real-world scientific problem with an ideal few-shot setting: there are many related low-data tasks convenient for knowledge transfer between each other. Furthermore, with cell images as a molecular representation, methods in the computer vision domain can be adapted for activity prediction. We believe the field of cell imaging/analysis would greatly benefit from these algorithmic innovations.

In this paper, we make two contributions:

- A data set for few-shot prediction of small molecule activity using cell microscopy images, which we named FSL-CP. The dataset is curated such that it is easy for future researchers to experiment their few-shot methods on.
- A benchmark of models encompassing different existing single-task, multi-task and meta-learning approaches on the dataset. This acts as both a diverse baseline for future algorithms, and a means to study the strength and weaknesses of different modelling paradigms.

## Methods

### FSL-CP: Few-Shot Learning Data Set with Cell Microscopy Images

The FSL-CP dataset comprises compounds in the intersection of ChEMBL^18^ version 31 and the Cell Painting^4^ public dataset. We provide an overview of the data construction process below (Figure 1), and the exact reproducible source code is available on GitHub, at https://github.com/czodrowskilab/FSL_CP_DataPrep.

**Figure 1:**
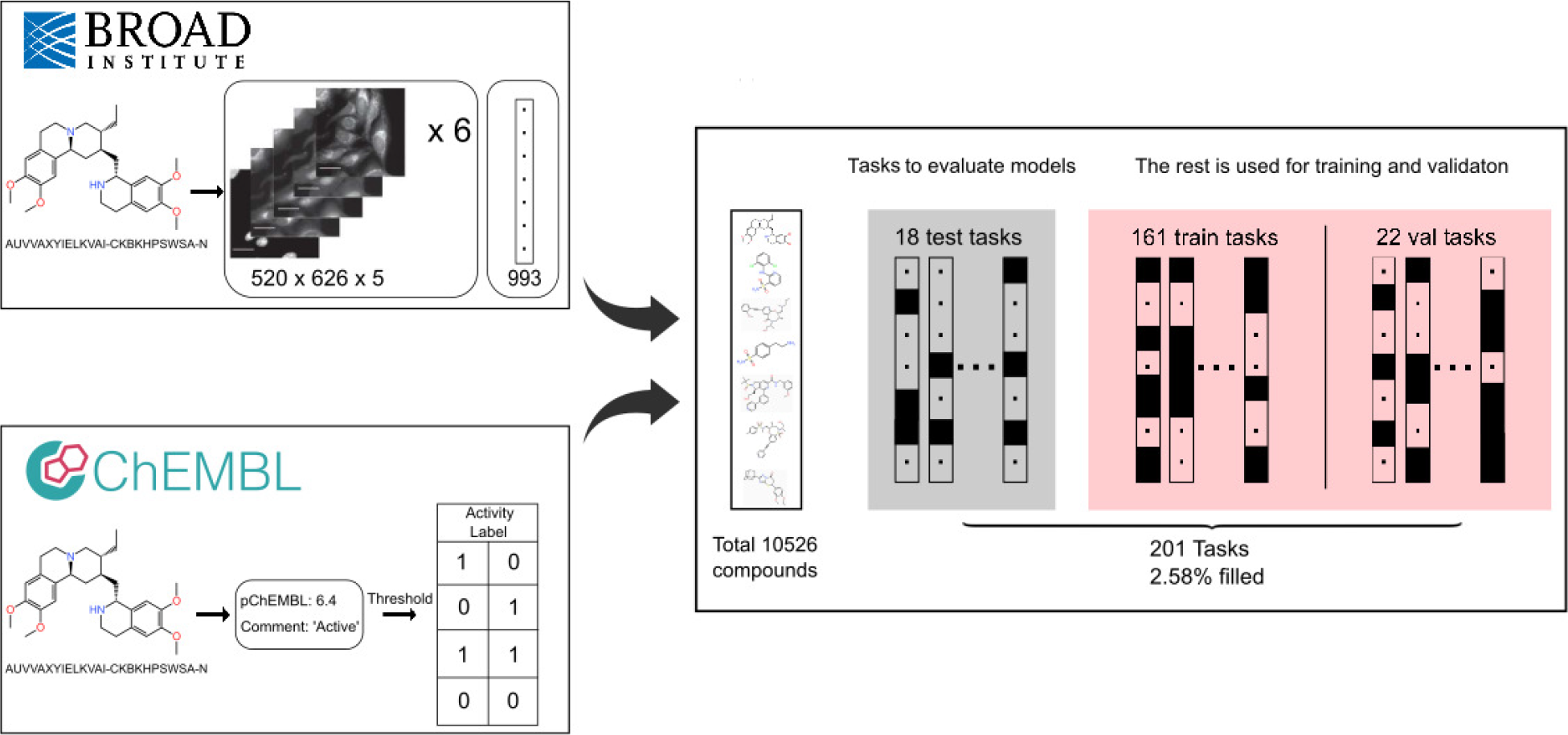
FSL-CP Data Curation and Processing. Cell painting images and features from CellProfiler ^19^ comes from Bray et al. ^4^ Each well is represented by six 520 × 696 × 5 images, and a feature vector of length 993. Small molecule activity labels are retrieved from assays in ChEMBL31. ^18^ Then a threshold procedure is applied to binarise the labels, producing different *tasks* from assays. The intersection of the two sources results in 10526 unique compounds, and 201 prediction tasks. 18 tasks are chosen for model evaluation based on a set of criteria, and the rest are for training and validation (referred to as *auxiliary tasks*.)

#### Original Cell Painting data set

Cell microscopy images come from from Bray et al.^4^ which contains 919,265 five-channel views, representing 30,616 compounds. In this Cell Painting protocol, U2OS cells have 8 major organelles and sub-compartments stained using a mixture of 6 fluorescent dyes, resulting in 5 different image channels. A CellProfiler^19^ pipeline is then used to extract 1,783 single-cell morphological features from those images.

#### Labelling the compounds

For this project we focus on small molecule activity assays (e.g. IC50 and EC50) available in the ChEMBL database. We query activity data for all the compounds in the original Cell Painting dataset using their InChiKey. We follow a similar data processing strategy as in Hofmarcher et al. ^20^ For each assay, both the *activity comments* from the experimenter and the *pChEMBL values* (numerical value for activity on a negative logarithmic scale) are retrieved. Duplicate labels are resolved either by averaging if they are pChEMBL values, or majority voting if they are activity comments. The pChEMBL values are restricted to only between 4 and 10, and the activity comments are also chosen to only be spelling variants of ‘Active’ and ‘Inactive’. The final modeling *tasks* is defined as an assay after being binarized, either with a threshold on the pChEMBL value, or based on the activity comments. For the pChEMBL values specifically, we use three thresholds for each assay: 5.5, 6.5, 7.5, which results in three separate modelling tasks. Lastly, we filter out to only allow tasks with at least 10 active and 10 inactive labelled compounds.

#### Processing the Cell Painting data

The images, as well as morphological features aggregated at well-level and metadata, can be found at the ‘Cell Image Library’. The five dye channels are concatenated along the third dimension, converted into 8 bits, and have their 0.0028% outlier bits removed.^20^ The images are further normalised prior to modelling. For the well-level morphological features, we remove columns that are highly correlated (correlation coefficient *>* 0.95), or have only one value. Finally, we standardize features by removing the mean and scaling to unit variance.

#### Features

At the end, each data point of FSL-CP corresponds to one well, represented by six 520×696×5 images (for six views in a well), and by a feature vector of length 993. We refer to them as CP images and CP features, respectively. SMILES strings and InChiKey are also provided, although for this study we only focus on the cell images and information which comes from them.

#### Deep Learning embedding

We also create an ‘enhanced’ set of CP features by concatenating the original CP features with embeddings from a ResNet50^21^ pretrained on ImageNet,^22^ akin to the method used in Schiff et al.^23^ This embedding provides the input vector with abstract high-level neural-network-based features. For each well, we run the six 520 × 696 × 5 views through the ResNet50 to generate six embeddings, which are then averaged to create one final embedding of length 1000. We tried different variants of ResNet and Inception, ^24,25^ before settling on ResNet50, which yield the best performance on our dataset despite being the simplest model. It should be noted that the length of the embedding can be further tuned to boost predictive performance.

##### Pretrain, validation and test splits

The models are evaluated on 18 tasks which we will call *test set D*_*test*_. The other 183 tasks, referred to as *auxiliary tasks*, are used for model pre-training. They are randomly splitted into *train set D*_*train*_ and *validation set D*_*val*_, consisting of 161 and 22 tasks, respectively. The test tasks are selected based on the following criteria:

- Tasks in *D*_*test*_ does not share the same targets as those in the *D*_*train*_ and *D*_*val*_, unless a target is unknown (denoted ‘unchecked’ on ChEMBL). This is to avoid the overlap of very similar tasks during training and inference.
- Test tasks must have over 96 datapoints, to enable model comparison for a range of support set sizes.
- Test tasks must have a ratio of active compound between 0.3 and 0.7, to avoid strongly imbalanced data affecting the model comparison.

##### Dataset statistics

FSL-CP contains 201 modelling tasks for 10526 unique compounds, with only 2.58% of the label matrix filled. Number of compounds for each task and their active ratio is visualised in (Figure 2A) and (Figure 2B), respectively. It is worth noting that the majority of the compounds has 4 replicates, as per the design of the cell painting assay. However, there are cases where there are fewer or more replicates (Figure 2C), potentially due to the omission of low-quality images and repeated purchase of some compounds.

**Figure 2:**
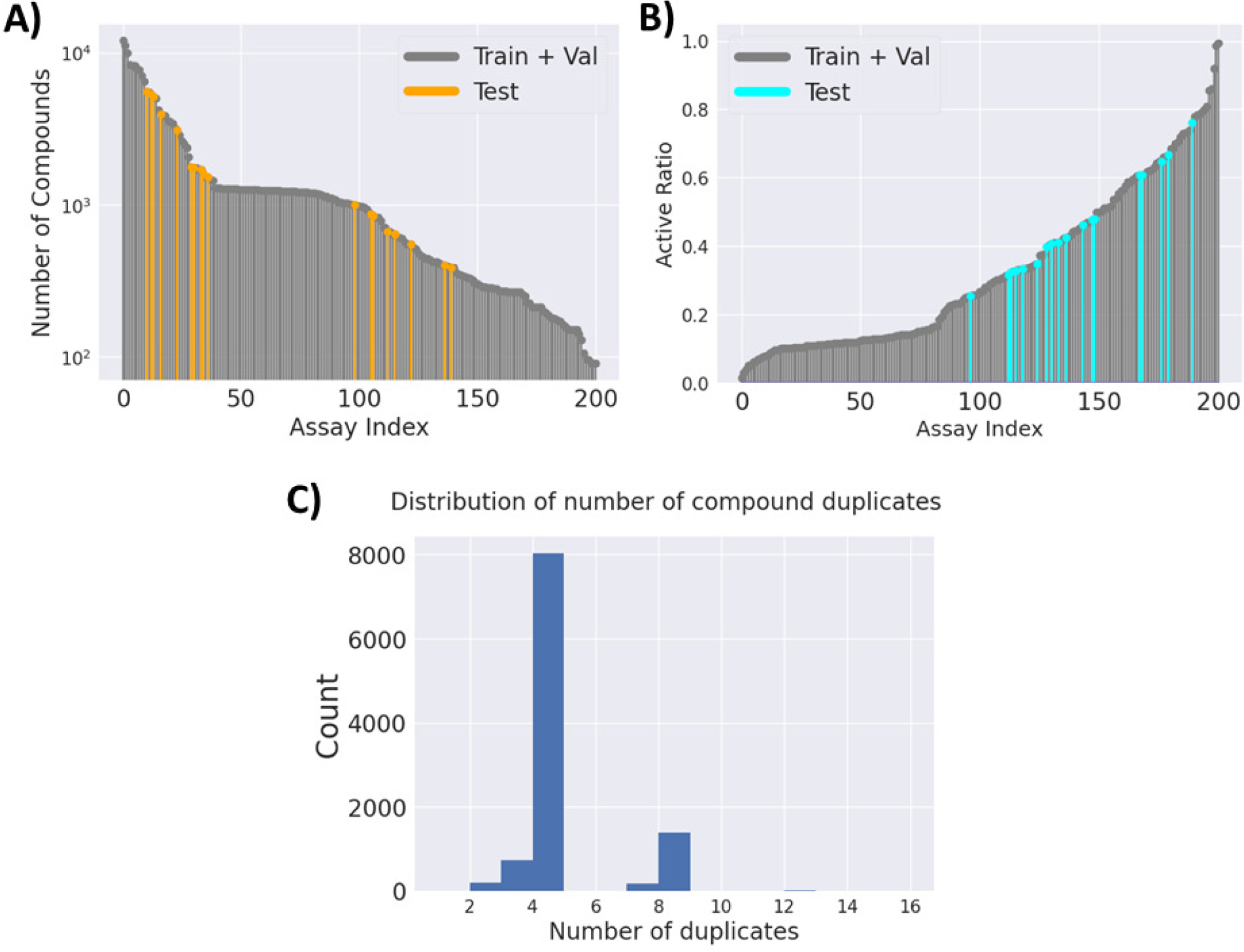
FSL-CP Data Statistics. (A) Number of compounds for every modelling task. (B) Ratio of active compounds for every modelling task. Test tasks (in turquoise) have their active ratio between 0.3 and 0.7. (C) Distribution of compound duplicates. Most have 4 duplicates as per experimental design, but there can be more or less duplicates, due to omission of low-quality images, or repeated purchases of compounds.

## Evaluation

### Few-shot prediction

In order to simulate the low-data setting when evaluating models, we sample from each task in a stratified manner a small number *M* of datapoints for the support set, and 32 datapoints for the query set. The models are then trained on a binary classification task on the support set, and evaluated on the query set. In literature, this sampled subset is called a *M* -samples 2-shot *episode*, where the ‘samples’ refer to available data (size of support set), and ‘shot’ refers to the number of classes to predict, which is two for binary classification.

For every test task, we report model performances averaged over 100 episodes in order to eliminate variations from sampling. Additionally, results are recorded over a range of support set sizes *M* : 8, 16, 32, 64, 96, to monitor how well models perform as size of available data increases.

### Metrics

We mainly report and discuss results using area under the receiver operating characteristic curve (AUROC). AUROC comes with many benefits, such as ranking predictions without a decision threshold, meaning predictions can be compared without needing to be rounded to 0 or 1. At the same time, the active ratio of tasks in *D*_*test*_ are not too imbalanced that they make AUROC unreliable.

In addition, results reported in F1 score, balanced accuracy, Cohen’s kappa, and ΔAUPRC^17^ can also be found in the Supplementary Information.

## Benchmark models

In this section, we provide a detailed description of different modelling paradigms for this particular few-shot problem. The code to all of the models and the training/inference scripts can be found at https://github.com/czodrowskilab/FSL_CP. As a naming convention, models with _**img** are trained directly on the images, _**cp** means they are trained on the original CP features, and _**cp+** means they are trained on the enhanced CP features.

### Single-task models

Traditionally, modelling of tasks in drug discovery is solely single-task, with models such as random forest or gradient boosting algorithms on top of fingerprints or curated phys-chem properties. ^26–29^ In these settings, auxiliary tasks are not used. Here, we mimic the same procedure by assessing the performance of logistic regression (LR), XGBoost, and a single-task fully-connected neural network (FNN), on both the original and enhanced CP features. For each prediction task of each model we run a randomised hyperparameter grid search using the library scikit-learn, ^30^ considering 10 hyperparameter configurations each run. We report results of the two best performing single-task models: LR on enhanced CP features (**logistic_cp+**) and FNN on original CP features (**singletask_cp**).

### Multi-task models

Multi-task models have been a staple in drug discovery field, being adopted by many academic and industry groups for various prediction tasks.^11,31,32^ These models consist of multiple ‘heads’, each specialised on one task, on top of a shared ‘trunk’. The trunk aims to learn a common representation across tasks, which allows it to learn knowledge transferable between tasks and improve performance of each one.

For our benchmark, the same FNN model as in the single-task case is used, but with a head of length 183 instead. Pretraining and validation are performed using 183 auxiliary tasks in *D*_*train*_ and *D*_*val*_ in a multi-task manner. Then the weights are frozen, and the head is replaced with a new one of length 1 for fine-tuning. During evaluation, for each episode, the same frozen model has its last layer fine-tuned using the support set, evaluated on the query set, and reverted back to the state before fine-tuned. We tried training on both sets of CP features but neither leads to drastic improvements over the other. We decided to report the result for the model trained on the original CP to reflect the methods from Simm et al. This model is denoted as **multitask_cp**.

### Meta-learning models

Inspired by human’s ability to learn certain tasks very quickly with prior knowledge, meta-learning methods aim to tackle the problem of adapting to new tasks efficiently with only a few training examples. The idea is still the same: pretrain to gain transferable knowledge to generalise to new tasks. But the meta-learning methods introduce the idea of *training in the same way as testing* .^33^ In particular, if we evaluate the model on *M* -samples 2-shot episodes, then we can mimic that setting during pretraining to encourage fast adaptation. That means during pretraining, we sample an episode from *D*_*train*_ the same way we sample from *D*_*test*_, and accumulate the loss from many episodes to update our models’ weights. This process is called *episodic training*.

One subclass of meta-learning is *metric-based method*, which tries to learn a distance function over data samples. For example, prototypical network ^12^ uses a backbone model to generate an embedding. Then classification is made using a k-means clustering based on the euclidean distance from the embedding to the cluster prototypes. Since the backbone for prototypical network can be any kind of embedding generator, we try 2 versions: a ResNet50 and an FNN backbone, which generate embeddings from CP images and CP features, respectively. These models are named **protonet_img, protonet_cp** and **protonet_cp+**.

*Optimisation-based* method is another subclass of meta-learning. This approach intends to make gradient-based optimisation converge within a small number of optimisation steps. MAML^13^ (Model-Agnostic Meta-Learning) achieves this by obtaining good weight initialisation through pretraining, so that fine-tuning to unseen tasks can be more efficient. Thanks to MAML working with any algorithm that uses gradient descent, we provide results of a ResNet50 and an FNN after being trained by MAML (denoted as **maml_img** and **maml_cp+**).

It is important to mention that unlike feature-based models, image-based models are highly computationally expensive to train. Hence to train these models and tune the hyperparameters in a reasonable time frame, only one out of six available views are used, plus random cropping and down-sizing of images are performed. With an 11GiB NVIDIA GeForce RTX 2080 Ti, the training and inference for one hyperparameter configuration takes between 1 and 2 weeks, depending on the model.

## Results

In order to compare performances of different methods benchmarked on FSL-CP, we plot the mean AUROC across 18 test tasks of each method at different support set sizes (Figure 3A). In addition, a paired Wilcoxon sign-rank test is performed for each pair of models as demonstrated in Table 1, with the alternative hypothesis that the column method outperforms the row method. These figures also provide insight into how performance of each model changes as the amount of available data increases.

**Table 1:** p-values of the one-sided Paired Wilcoxon Sign-rank Test^*a*^ with Alternative Hypothesis being the method in the left column outperforms the method in the upper row. Entries left blank indicate a p-value greater or equal to 9.99e-01.

|  | protonet_cp | multitask_cp | singletask_cp | maml_cp+ | logistic_cp+ | maml_img | protonet_img |
| --- | --- | --- | --- | --- | --- | --- | --- |
| protonet_cp+ | 3.60e-01 | 7.89e-01 | 1.56e-01 | 3.79e-01 | **1.05e-03** | **3.09e-03** | **5.38e-04** |
| protonet_cp |  | 8.90e-01 | 2.74e-01 | 6.12e-01 | **1.79e-03** | **5.23e-03** | **4.52e-04** |
| multitask_cp |  |  | 3.96e-02 | 3.04e-01 | **2.69e-04** | **1.11e-03** | **1.43e-04** |
| singletask_cp |  |  |  | 6.03e-01 | **3.09e-03** | 8.05e-02 | **1.21e-03** |
| maml_cp+ |  |  |  |  | 8.84e-02 | 1.92e-02 | 2.74e-02 |
| logistic_cp+ |  |  |  |  |  | 6.42e-01 | 9.16e-03 |
| maml_img |  |  |  |  |  |  | 3.47e-02 |

b) Support set size 16
|  | protonet_cp | multitask_cp | singletask_cp | maml_cp+ | logistic_cp+ | maml_img | protonet_img |
| --- | --- | --- | --- | --- | --- | --- | --- |
| protonet_cp+ | 1.45e-02 | 5.42e-02 | **9.36e-04** | **2.77e-03** | **2.49e-04** | **5.34e-05** | **7.63e-06** |
| protonet_cp |  | 5.41e-01 | 1.08e-02 | 2.21e-02 | **8.53e-04** | **5.35e-04** | **2.63e-04** |
| multitask_cp |  |  | **5.22e-04** | 1.13e-02 | **2.08e-04** | **9.54e-05** | **1.46e-04** |
| singletask_cp |  |  |  | 6.51e-01 | **1.68e-03** | 3.49e-02 | **3.43e-04** |
| maml_cp+ |  |  |  |  | 2.01e-02 | **2.98e-03** | **1.03e-03** |
| logistic_cp+ |  |  |  |  |  | 2.48e-01 | **1.50e-03** |
| maml_img |  |  |  |  |  |  | 1.27e-01 |

c) Support set size 32
|  | protonet_cp | multitask_cp | singletask_cp | maml_cp+ | logistic_cp+ | maml_img | protonet_img |
| --- | --- | --- | --- | --- | --- | --- | --- |
| protonet_cp+ | 8.63e-02 | **7.97e-03** | **3.83e-04** | **1.42e-04** | **1.46e-04** | **3.81e-06** | **3.81e-06** |
| protonet_cp |  | 6.49e-02 | **1.46e-03** | **2.67e-04** | **2.67e-05** | **1.74e-04** | **1.14e-05** |
| multitask_cp |  |  | **4.03e-04** | **4.56e-03** | **3.81e-06** | **3.81e-06** | **3.81e-06** |
| singletask_cp |  |  |  | 3.61e-01 | **2.80e-03** | **1.22e-03** | **2.56e-04** |
| maml_cp+ |  |  |  |  | **3.69e-02** | **1.65e-03** | **5.23e-04** |
| logistic_cp+ |  |  |  |  |  | **7.97e-03** | **1.36e-04** |
| maml_img |  |  |  |  |  |  | 2.55e-01 |

d) Support set size 64
|  | protonet_cp | multitask_cp | singletask_cp | maml_cp+ | logistic_cp+ | maml_img | protonet_img |
| --- | --- | --- | --- | --- | --- | --- | --- |
| protonet_cp+ | 2.62e-01 | 3.43e-01 | **6.97e-03** | **4.40e-04** | **7.63e-06** | **3.81e-06** | **3.81e-06** |
| protonet_cp |  | 3.88e-01 | 1.46e-02 | **6.05e-04** | **1.91e-05** | **1.46e-04** | **7.63e-06** |
| multitask_cp |  |  | **2.07e-04** | **2.47e-04** | **3.81e-06** | **3.81e-06** | **3.81e-06** |
| singletask_cp |  |  |  | 9.31e-02 | **1.43e-04** | **3.81e-06** | **3.81e-06** |
| maml_cp+ |  |  |  |  | **4.10e-03** | **1.14e-05** | **1.14e-05** |
| logistic_cp+ |  |  |  |  |  | **2.02e-03** | **2.11e-04** |
| maml_img |  |  |  |  |  |  | 4.18e-01 |

e) Support set size 96
|  | protonet_cp | multitask_cp | singletask_cp | maml_cp+ | logistic_cp+ | maml_img | protonet_img |
| --- | --- | --- | --- | --- | --- | --- | --- |
| protonet_cp+ | 6.14e-01 | 2.84e-01 | 7.73e-02 | **7.25e-05** | **3.81e-06** | **3.81e-06** | **3.81e-06** |
| protonet_cp |  | 2.09e-01 | 6.44e-02 | **2.29e-04** | **2.16e-04** | **3.81e-06** | **3.81e-06** |
| multitask_cp |  |  | 4.00e-02 | **1.45e-04** | **3.81e-06** | **3.81e-06** | **3.81e-06** |
| singletask_cp |  |  |  | **3.36e-04** | **1.44e-04** | **3.81e-06** | **3.81e-06** |
| maml_cp+ |  |  |  |  | 1.85e-01 | **1.14e-05** | **4.44e-04** |
| logistic_cp+ |  |  |  |  |  | **3.21e-04** | **3.81e-06** |
| maml_img |  |  |  |  |  |  | 2.33e-01 |

<sup>a</sup> The test compares mean AUROC (over 100 episodes) of models in 18 test tasks. Marked in bold are significant p-values at $\alpha = 0.01$ .

**Figure 3:**
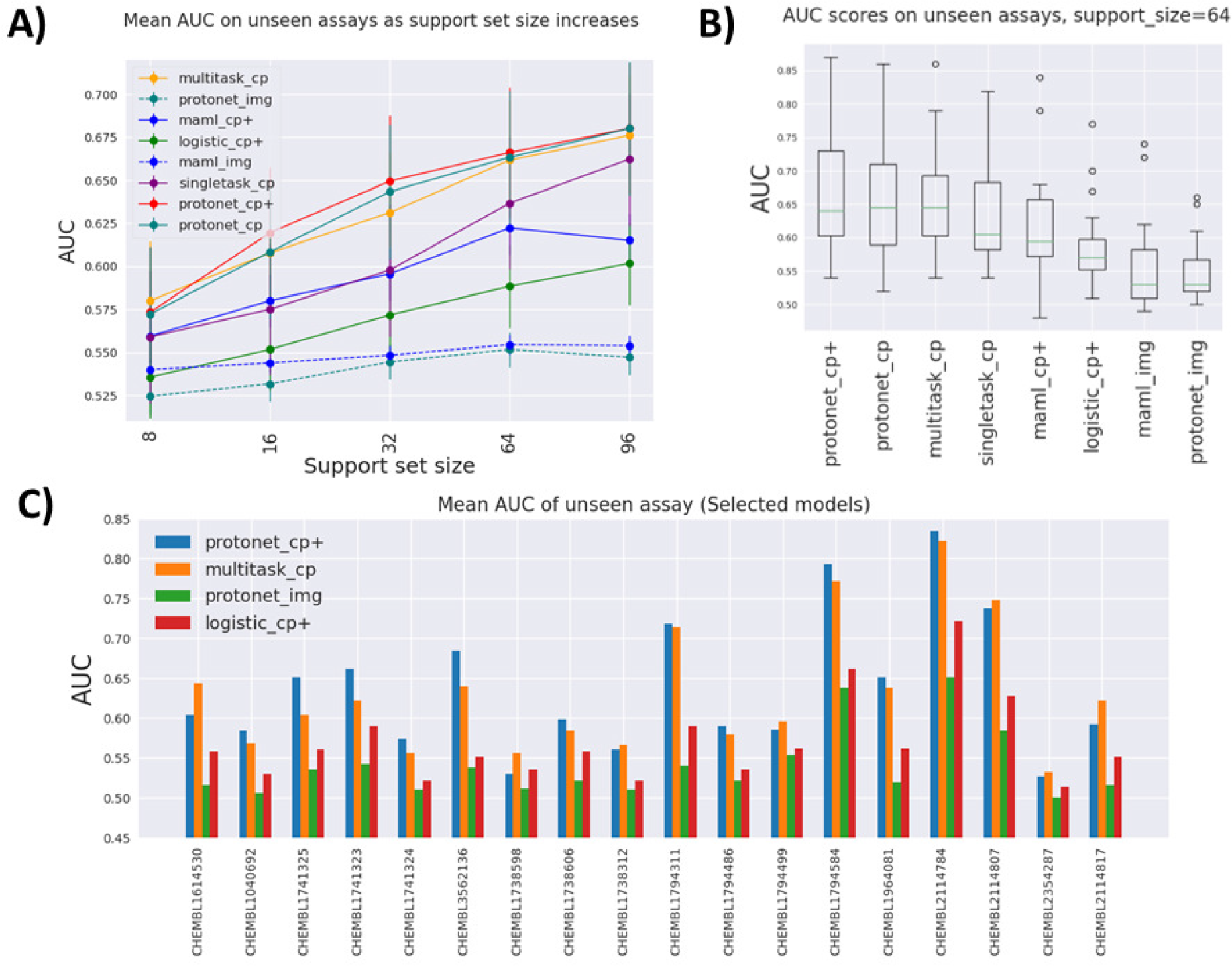
Comparison of different models benchmarked on FSL-CP. Figure A): Mean AUROC on test tasks as support set size increases. As there are more data available, other methods start to catch up to meta-learning models. Figure B): Distribution of AUROC across all test tasks at support set size 64. The best models tend to have larger AUROC variance. Figure C): Mean AUROC of selected models for each task across all support set sizes. For most tasks, pretraning on auxiliary tasks leads to an improvement over single-task models. However, for a few tasks this is not the case.

Figure 3A indicates that the best performing models overall are variants of prototypical network **protonet_cp+** and **protonet_cp**, followed closely by **multitask_cp**. Although according to the Wilcoxon signed-rank test, **protonet_cp+** only outperforms **multitask_cp** for medium-sized dataset (support set size 32). There is not sufficient evidence rejecting the null hypothesis that they perform equally well at other support set sizes with *α* = 0.01. We note that **multitask_cp** even slightly outperforms **protonet_cp+** and **protonet_cp** at support set size 8.

The best single-task model **singletask_cp** is surprisingly powerful, being able to catch up with **maml_cp+** at lower support set size, and outperforms it by a wide margin at large support set size. For these single-task models with no pretraining, the availability of more data in the support set can lead to dramatic improvements in performance. It is highly likely that their performances will keep improving, and eventually might overtake other methods beyond support set size 96. On the contrary, improvements in AUROC scores of metalearning methods slows down at higher support set size, and even drops as in the case of **maml_cp+**. However, it is worth noting that at support set size 96, some test tasks are excluded in the evaluation process due to insufficient data points. Plus, less tasks in *D*_*train*_ and *D*_*val*_ are included in the pretraining for meta-learning models at high support set size, due to the fact that there may not be enough datapoints to sample for episodic training. All of these factors can affect meta-learning methods’ performance at higher data setting.

Image-based models such as **protonet_img** and **maml_img** substantially under-perform compared to other feature-based methods, likely because only one view out of six is used, and down-sizing of images of fairly small cells leads to drastic information loss.

We also try to leverage deep-learning-based features by concatenating the original CP features with a ResNet50 embedding of size 1000. While in some cases it does lead to higher AUROC, as evidenced by the fact that many models use the enhanced feature, the improvements are somewhat minute. For example, when comparing **protonet_cp+** against **protonet_cp**, Table 1 shows insufficient evidence of improvement across tasks, and in Figure 3A, the additional features lead to only small improvements at support set sizes 16, 32 and 64. However, this still poses an interesting question for future research: how meaningful embedding from cell images can be produced using deep learning methods.

Better performing models have the larger spread of AUROC across test tasks, as shown in Figure 3B, indicating model performances are fairly dependent on tasks. This is further demonstrated in Figure 3C, where some tasks (e.g. CHEMBL2114784) consistently show high AUROC across model, and some (e.g. CHEMBL2354287) are not predictive at all. Additionally, for some tasks **protonet_cp+** is the best method, but in a few other cases **multitask_cp** or **singletask_cp** is the better method.

Figure 3C also gives insight on how much pretraining on auxiliary tasks benefits prediction. Again, this is highly task-dependent. Some tasks benefit greatly from pretraining (CHEMBL3562136, CHEMBL2114807), as seen from the improvements of the two pretraining models over the single-task models. However, pretraining can offer no improvement, or even be detrimental in tasks such as CHEMBL1738598 and CHEMBL1738312.

## Discussion

We have presented FSL-CP, a dataset for small molecule activity few-shot prediction using cell microscopy images. This few-shot challenge mimics a screening process in early stage drug discovery, where the aim is to identify potent compounds targeting a specific protein from high-content cell images with little data. Previous efforts have been made to benchmark few-shot methods on molecules as graph-structured data.^17^ But in machine learning, the primary focus of few-shot learning has been in the computer vision and natural language processing domain. The fact that our dataset uses cell images as a molecular representation opens up opportunities to adapt state-of-the-art ideas from computer vision to enhance modeling.

This dataset allows us to establish benchmarks that compare the performances of different few-shot learning paradigms. Our result indicates feature-based prototypical network and multitask FNN pretrained on auxiliary tasks generally preforms well across all support set sizes. We also observe improvement in performance slowing down for meta-learning methods at high support set size, in contrast to single-task methods, which greatly benefits from the availability of more available data. However, more labelled compounds and their cell painting data are needed in order to accurately point out whether eventually single-task models outperforms pretrained models, and if yes, at what support set size.

Image-based models underperforms on our benchmark, and the fact that each datapoint consists of six high-definition five-channel images makes it tremendously computationally expensive to train. We had to use only one randomly cropped, downsized image to train the models in a reasonable time-frame with our infrastructure, and this leads to high information loss. Training on full-resolution cell images has been shown to offer better performance than on CP features in some settings. ^20^ However, in realistic drug discovery projects, larger-size images are used, and there typically are more compounds and more prediction tasks. These make pretraining on full-resolution images difficult, especially if the model needs to be regularly retrained.

A less expensive way to leverage the power of computer vision is to enhance the CP feature with an embedding out of an image using a pretrained model such as ResNet or Inception. We tried a simple approach with a ResNet50 as an embedding generator, which yielded small improvements. Since most vision models are pretrained on ImageNet, this suggests there is some transferable knowledge obtained from training on a big unrelated image database, but not enough to make a significant improvement. We expect that a more informative embedding can be achieved by pretraining the embedding generator end-to-end on cell images with a more relevant pretraining task, such as multi-task or contrastive learning.^34,35^

The benchmark also provides insight on how effective transferring knowledge from pretraining models on auxiliary tasks is, to new tasks. Mostly, new tasks benefit from such a pretraining scheme, but the degree to which different tasks improve varies. To what degree a new task benefits from pretraining is still an open question for research. As a general observation, it seems already predictive tasks tend to benefit more from pretraining. Companies aiming to pretrain their models on auxiliary tasks can use a combination of tasks from public sources as well as their own database to benefit from as much data as possible.

FSL-CP offers a promising and interesting research question, though not one without unique challenges of its own. Through this study, we hope to encourage further research in few-shot and computer vision methods in the domain of cell imaging. The code for generating the dataset, model training, inference, and tutorials are publicly available on GitHub.

## Acknowledgement

The first author’s research position is funded by the the European Union’s Horizon 2020 research and innovation program under the Marie Sklodowska-Curie Innovative Training Network - European Industrial Doctorate grant agreement No. 956832, “Advanced machine learning for Innovative Drug Discovery”. The authors thank Steffen Jaensch, Lorena G.A. Freitas for proofreading the manuscript, and others from Janssen Pharmaceutical for providing feedback for the project. In addition, we thank Axel Pahl and Jose Manuel-Gally from the Max-Planck Institute in Dortmund, and other members of the Czodrowski lab for the fruitful discussions.

## References

(1) Moffat, J. G., Rudolph, J., Bailey, D. Phenotypic screening in cancer drug discovery - past, present and future. Nat. Rev. Drug Discov. 2014, 13, 588–602.

(2) Johannessen, C. M., Clemons, P. A., Wagner, B. K. Integrating phenotypic small-molecule profiling and human genetics: the next phase in drug discovery. Trends Genet. 2015, 31, 16–23.

(3) Herman, D. et al. Leveraging Cell Painting Images to Expand the Applicability Domain and Actively Improve Deep Learning Quantitative Structure–Activity Relationship Models. Chemical Research in Toxicology 2023, 36, 1028–1036.

(4) Bray, M.-A. et al. A dataset of images and morphological profiles of 30 000 smallmolecule treatments using the Cell Painting assay. GigaScience 2017, 6, giw014.

(5) Caicedo, J. C., Singh, S., Carpenter, A. E. Applications in image-based profiling of perturbations. Current Opinion in Biotechnology 2016, 39, 134–142.

(6) Reisen, F., Sauty de Chalon, A., Pfeifer, M., Zhang, X., Gabriel, D., Selzer, P. Linking phenotypes and modes of action through high-content screen fingerprints. Assay Drug Dev. Technol. 2015, 13, 415–427.

(7) Young, D. W., Bender, A., Hoyt, J., McWhinnie, E., Chirn, G.-W., Tao, C. Y., Tallarico, J. A., Labow, M., Jenkins, J. L., Mitchison, T. J., Feng, Y. Integrating highcontent screening and ligand-target prediction to identify mechanism of action. Nat. Chem. Biol. 2008, 4, 59–68.

(8) Ljosa, V., Caie, P. D., Ter Horst, R., Sokolnicki, K. L., Jenkins, E. L., Daya, S., Roberts, M. E., Jones, T. R., Singh, S., Genovesio, A., Clemons, P. A., Carragher, N. O., Carpenter, A. E. Comparison of methods for image-based profiling of cellular morphological responses to small-molecule treatment. J. Biomol. Screen. 2013, 18, 1321–1329.

(9) Collinet, C., Stöter, M., Bradshaw, C. R., Samusik, N., Rink, J. C., Kenski, D., Habermann, B., Buchholz, F., Henschel, R., Mueller, M. S., Nagel, W. E., Fava, E., Kalaidzidis, Y., Zerial, M. Systems survey of endocytosis by multiparametric image analysis. Nature 2010, 464, 243–249.

(10) Fuchs, F., Pau, G., Kranz, D., Sklyar, O., Budjan, C., Steinbrink, S., Horn, T., Pedal, A., Huber, W., Boutros, M. Clustering phenotype populations by genome-wide RNAi and multiparametric imaging. Mol. Syst. Biol. 2010, 6, 370.

(11) Simm, J. et al. Repurposing High-Throughput Image Assays Enables Biological Activity Prediction for Drug Discovery. Cell Chemical Biology 2018, 25, 611–618.e3.

(12) Snell, J., Swersky, K., Zemel, R. S. Prototypical Networks for Few-shot Learning. 2017.

(13) Finn, C., Abbeel, P., Levine, S. Model-Agnostic Meta-Learning for Fast Adaptation of Deep Networks. 2017.

(14) Brown, T. B. et al. Language Models are Few-Shot Learners. 2020.

(15) Geng, R., Li, B., Li, Y., Zhu, X., Jian, P., Sun, J. Induction Networks for Few-Shot Text Classification. 2019.

(16) Vinyals, O., Blundell, C., Lillicrap, T., Kavukcuoglu, K., Wierstra, D. Matching Networks for One Shot Learning. 2017.

(17) Stanley, M., Bronskill, J. F., Maziarz, K., Misztela, H., Lanini, J., Segler, M., Schneider, N., Brockschmidt, M. FS-Mol: A Few-Shot Learning Dataset of Molecules. NeurIPS 2021 Track Datasets and Benchmarks 2021,

(18) Mendez, D. et al. ChEMBL: towards direct deposition of bioassay data. Nucleic Acids Research 2018, 47, D930–D940.

(19) Carpenter, A., Jones, T., Lamprecht, M., Clarke, C., Kang, I., Friman, O., Guertin, D., Chang, J., Lindquist, R., Moffat, J., Golland, P., Sabatini, D. CellProfiler: Image analysis software for identifying and quantifying cell phenotypes. Genome biology 2006, 7, R100.

(20) Hofmarcher, M., Rumetshofer, E., Clevert, D.-A., Hochreiter, S., Klambauer, G. Accurate Prediction of Biological Assays with High-Throughput Microscopy Images and Convolutional Networks. Journal of Chemical Information and Modeling 2019, 59, 1163–1171.

(21) He, K., Zhang, X., Ren, S., Sun, J. Deep Residual Learning for Image Recognition. 2015.

(22) Deng, J., Dong, W., Socher, R., Li, L.-J., Li, K., Fei-Fei, L. ImageNet: A Large-Scale Hierarchical Image Database. CVPR09. 2009.

(23) Schiff, L. et al. Integrating deep learning and unbiased automated high-content screening to identify complex disease signatures in human fibroblasts. Nature Communications 2022, 13, 1590.

(24) Szegedy, C., Liu, W., Jia, Y., Sermanet, P., Reed, S., Anguelov, D., Erhan, D., Vanhoucke, V., Rabinovich, A. Going Deeper with Convolutions. 2014.

(25) Szegedy, C., Vanhoucke, V., Ioffe, S., Shlens, J., Wojna, Z. Rethinking the Inception Architecture for Computer Vision. 2015.

(26) Rogers, D., Hahn, M. Extended-Connectivity Fingerprints. Journal of Chemical Information and Modeling 2010, 50, 742–754, PMID: 20426451.

(27) Mauri, A., Consonni, V., Pavan, M., Todeschini, R. DRAGON software: An easy approach to molecular descriptor calculations. MATCH Communications in Mathematical and in Computer Chemistry 2006, 56, 237–248.

(28) Soufan, O., Ba-alawi, W., Magana-Mora, A., Essack, M., Bajic, V. B. DPubChem: a web tool for QSAR modeling and high-throughput virtual screening. Scientific Reports 2018, 8, 9110.

(29) Butina, D., Segall, M. D., Frankcombe, K. Predicting ADME properties in silico: methods and models. Drug Discov. Today 2002, 7, S83–8.

(30) Pedregosa, F. et al. Scikit-learn: Machine Learning in Python. Journal of Machine Learning Research 2011, 12, 2825–2830.

(31) Mayr, A., Klambauer, G., Unterthiner, T., Hochreiter, S. DeepTox: Toxicity Prediction using Deep Learning. Frontiers in Environmental Science 2016, 3 .

(32) Hu, W., Liu, B., Gomes, J., Zitnik, M., Liang, P., Pande, V., Leskovec, J. Strategies for Pre-training Graph Neural Networks. 2020.

(33) Weng, L. Meta-Learning: Learning to Learn Fast. lilianweng.github.io 2018, Accessed: 2023-04-26.

(34) Chaitanya, K., Erdil, E., Karani, N., Konukoglu, E. Contrastive learning of global and local features for medical image segmentation with limited annotations. 2020.

(35) Radford, A., Kim, J. W., Hallacy, C., Ramesh, A., Goh, G., Agarwal, S., Sastry, G., Askell, A., Mishkin, P., Clark, J., Krueger, G., Sutskever, I. Learning Transferable Visual Models From Natural Language Supervision. 2021.

