## Supplementary Information for "FSL-CP: A Benchmark for Small Molecule Activity Few-Shot Prediction using Cell Microscopy Images"

### List of Tables

1 **Test tasks**: Details of the assays in the test set and their targets in ChEMBL. 4

This document provides supplementary information for the publication "FSL-CP: A Benchmark for Small Molecule Activity Few-Shot Prediction using Cell Microscopy Images". We report details about the 201 tasks of the dataset, and additional performance metrics.

The dataset, model codes and performances of all models (including those not reported in the publication) are all publicly available on Github:

`https://github.com/czodrowskilab/FSL_CP`.

Table 1: **Test tasks**: Details of the assays in the test set and their targets in ChEMBL.

| TASK_ID | assay_chembl_id | target_chembl_id | target_type |
|---|---|---|---|
| 688267 | CHEMBL1614530 | CHEMBL2026 | SINGLE PROTEIN |
| 600886 | CHEMBL1040692 | CHEMBL364 | ORGANISM |
| 737826 | CHEMBL1741325 | CHEMBL3397 | SINGLE PROTEIN |
| 737824_1 | CHEMBL1741323 | CHEMBL3622 | SINGLE PROTEIN |
| 737825 | CHEMBL1741324 | CHEMBL340 | SINGLE PROTEIN |
| 1495405 | CHEMBL3562136 | CHEMBL612545 | UNCHECKED |
| 737053 | CHEMBL1738598 | CHEMBL612545 | UNCHECKED |
| 737400 | CHEMBL1738606 | CHEMBL5501 | SINGLE PROTEIN |
| 736947 | CHEMBL1738312 | CHEMBL1741220 | SINGLE PROTEIN |
| 752347 | CHEMBL1794311 | CHEMBL1977 | SINGLE PROTEIN |
| 752496 | CHEMBL1794486 | CHEMBL5027 | SINGLE PROTEIN |
| 752509 | CHEMBL1794499 | CHEMBL1795091 | SINGLE PROTEIN |
| 752594 | CHEMBL1794584 | CHEMBL1293258 | SINGLE PROTEIN |
| 809095 | CHEMBL1964081 | CHEMBL2007624 | SINGLE PROTEIN |
| 845173 | CHEMBL2114784 | CHEMBL1795085 | SINGLE PROTEIN |
| 845196 | CHEMBL2114807 | CHEMBL5990 | SINGLE PROTEIN |
| 954338 | CHEMBL2354287 | CHEMBL2362981 | SINGLE PROTEIN |
| 845206 | CHEMBL2114817 | CHEMBL4377 | SINGLE PROTEIN |

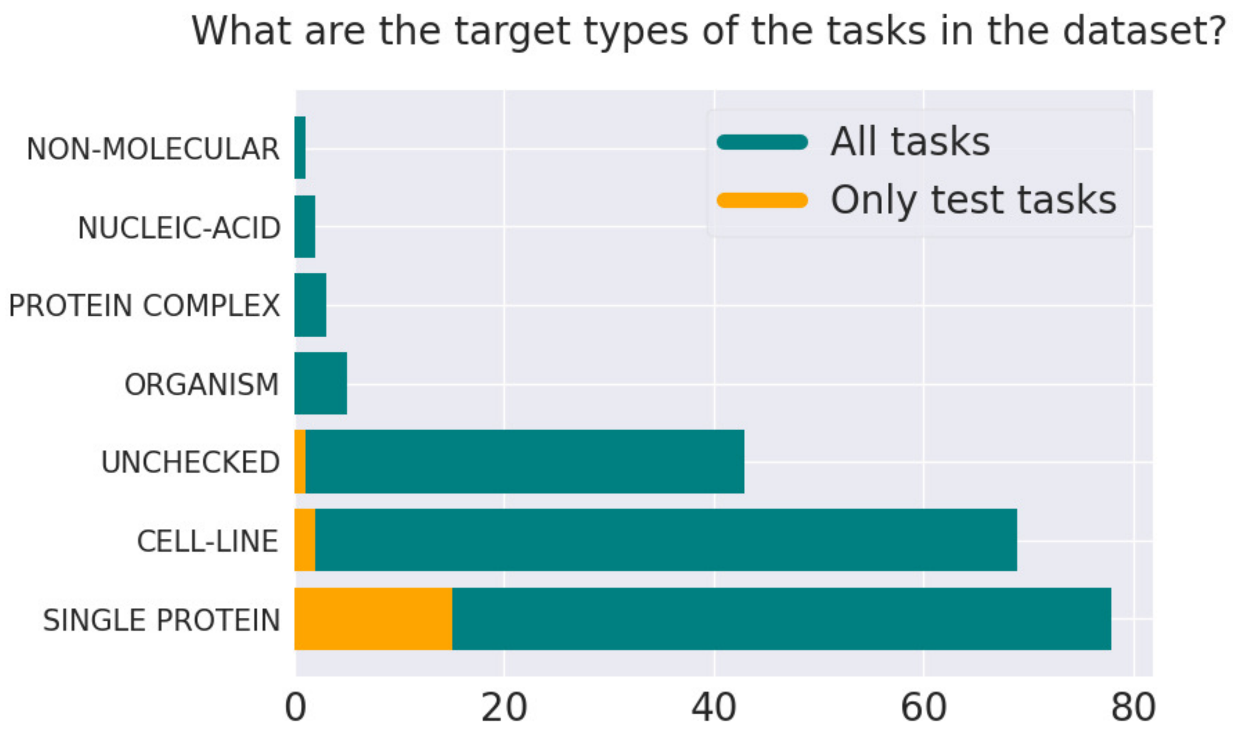

Figure 1: **What are the target types of the tasks in the dataset?** The majority are single proteins and cell lines, both in the entire set of tasks and the test tasks only.

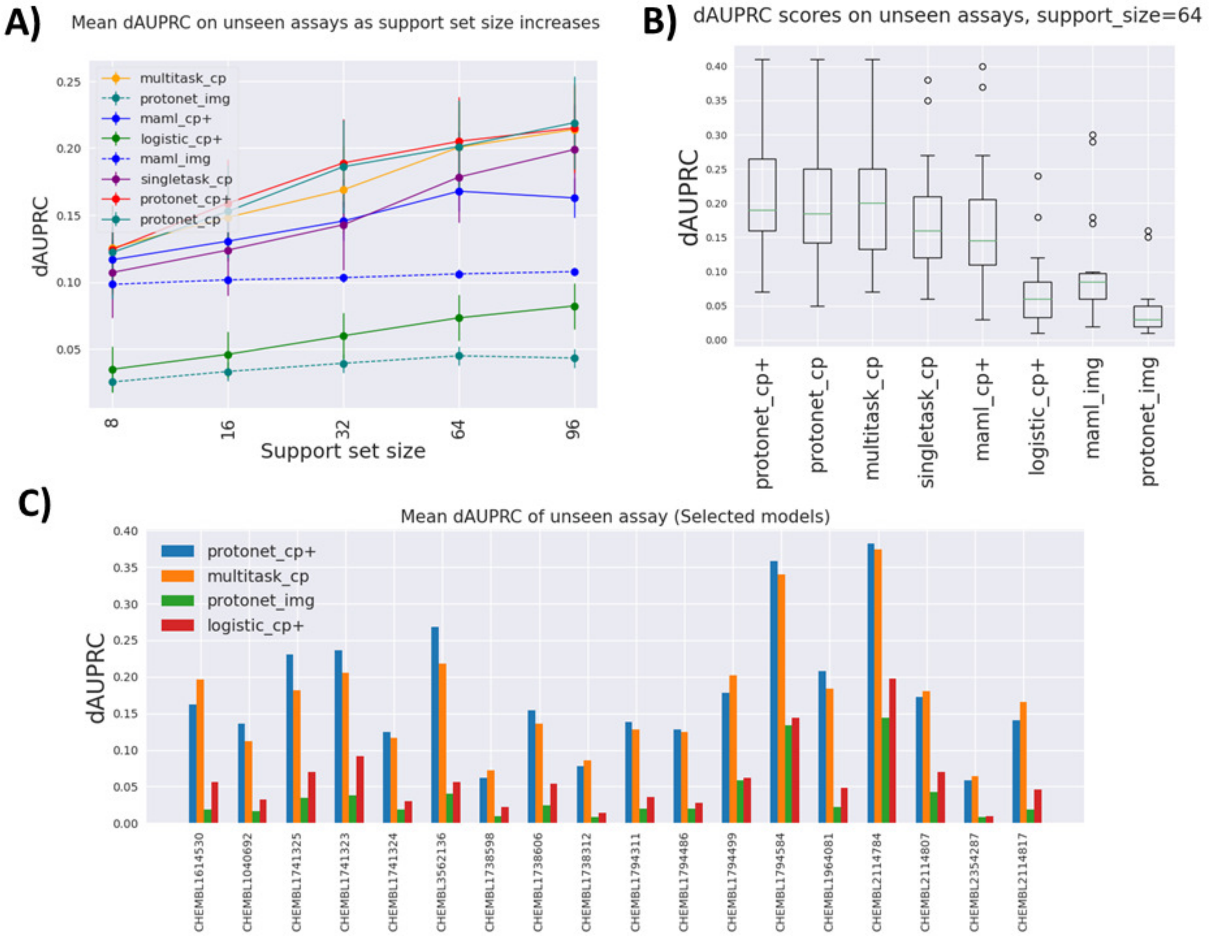

Figure 2: **Result reported using dAUPRC.** Figure A): Mean dAUPRC on test tasks as support set size increases. Figure B): Distribution of dAUPRC across all test tasks at support set size 64. Figure C): Mean dAUPRC of selected models for each task across all support set sizes.

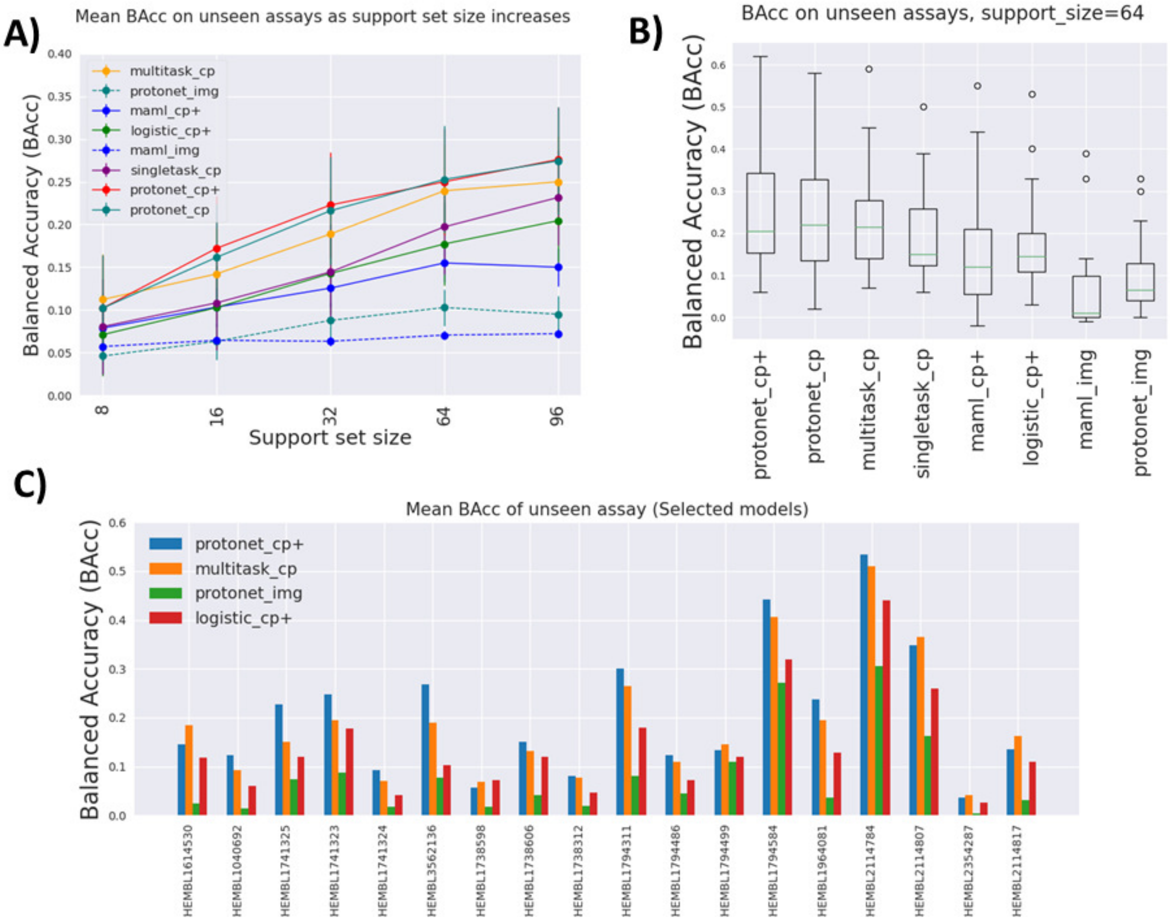

Figure 3: **Result reported using Balanced Accuracy (BAcc).** Figure A): Mean BAcc on test tasks as support set size increases. Figure B): Distribution of BAcc across all test tasks at support set size 64. Figure C): Mean BAcc of selected models for each task across all support set sizes.

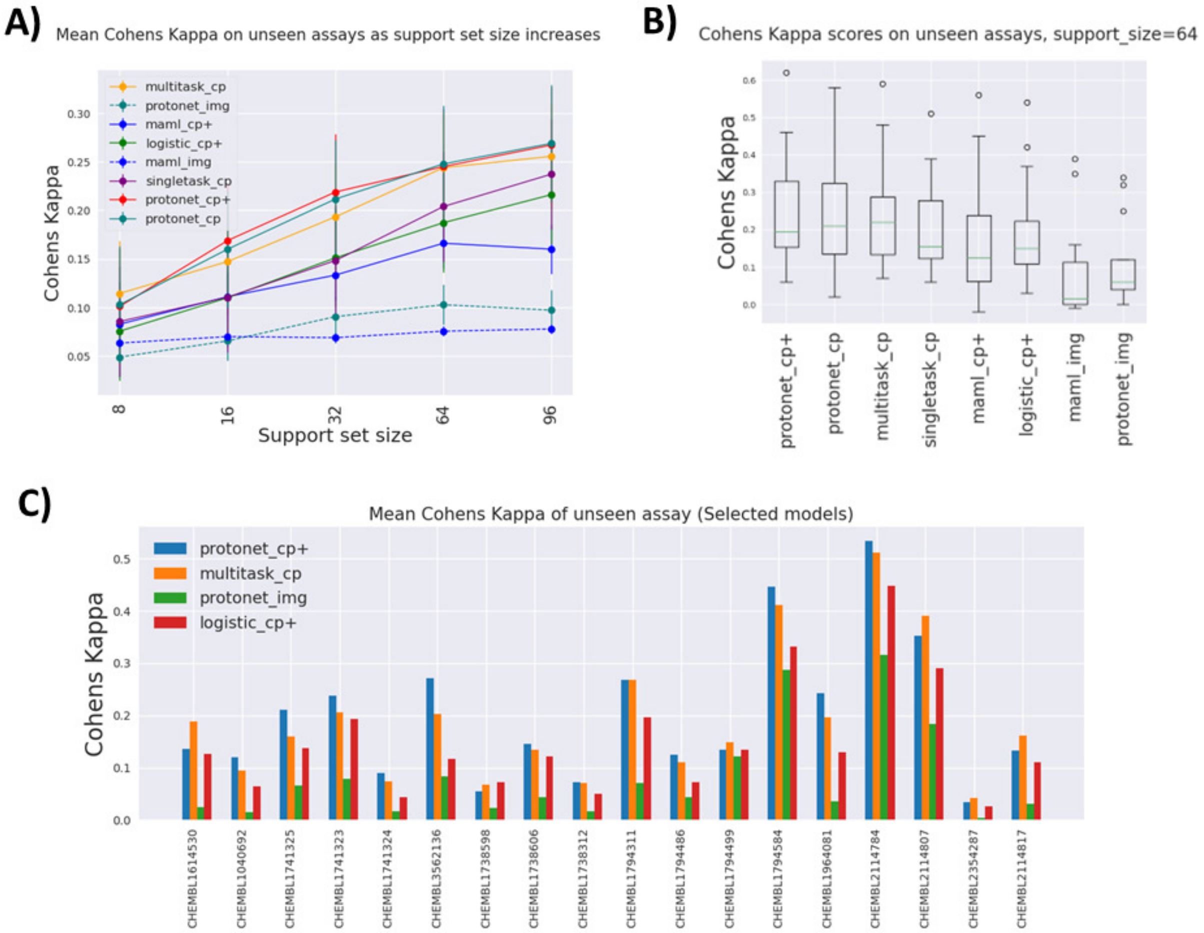

Figure 4: **Result reported using Cohens Kappa.** Figure A): Mean Cohens Kappa on test tasks as support set size increases. Figure B): Distribution of Cohens Kappa across all test tasks at support set size 64. Figure C): Mean Cohens Kappa of selected models for each task across all support set sizes.

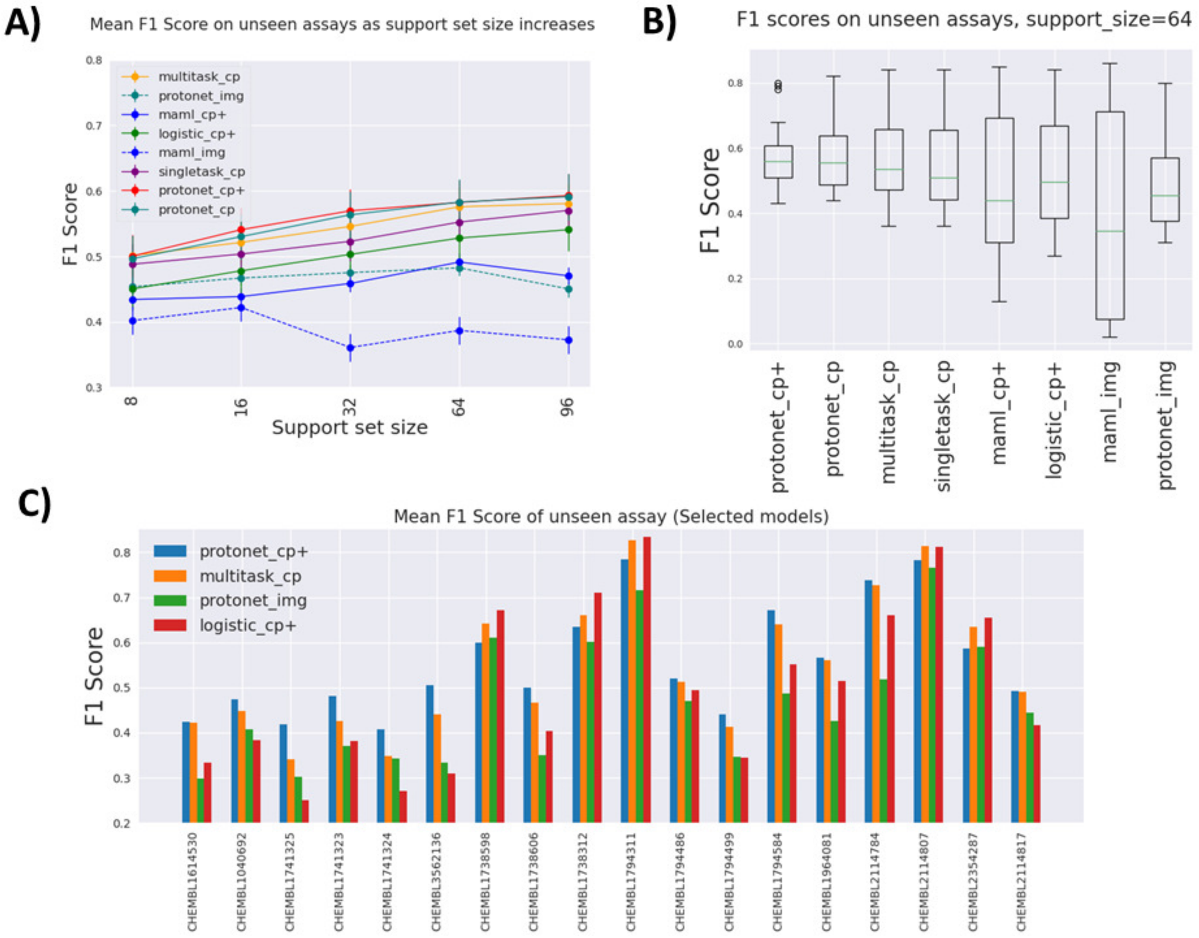

Figure 5: **Result reported using F1 Score.** Figure A): Mean F1 score on test tasks as support set size increases. Figure B): Distribution of F1 score across all test tasks at support set size 64. Figure C): Mean F1 score of selected models for each task across all support set sizes.
